# DELPHAI, AI Agent for Predicting Drug Response and Resistance

**DOI:** 10.1101/2025.09.12.675826

**Authors:** Tianping Peng, Hui Wu, Haikun Liu, Xian Zhang

**Affiliations:** aiPTO TechBio, Aeschengraben 29, 4051, Basel, Switzerland; Division of Molecular Neurogenetics, German Cancer Research Center (DKFZ), Im Neuenheimer Feld 581, 69120, Heidelberg, Germany

## Abstract

Patient-derived organoids preserve critical tumor features and drug sensitivity patterns that mirror patient clinical responses, enabling single-cell RNA sequencing analysis of drug responses. Analyzing these perturbation data presents significant computational challenges in predicting cellular responses while maintaining biological interpretability. We developed DELPHAI (Deep ExplainabLe Predictive Human-organoid based AI), an AI agent that integrates single-cell perturbation prediction with mechanistic analysis using large language models. We designed a comprehensive benchmarking framework evaluating methods in both computational embedding and reconstructed gene expression spaces. Applied to glioblastoma organoids treated with temozolomide, optimal transport combined with principal component analysis outperformed baseline methods in capturing population dynamics. DELPHAI correctly identified DNA alkylation as the mechanism of action without prior drug knowledge and recommended combination therapies aligning with clinical trials. These results demonstrate DELPHAPs ability to translate single-cell perturbation data into actionable therapeutic insights, representing a significant advance toward Al-driven precision medicine in cancer treatment.

## Introduction

Patient-derived organoids have emerged as transformative models for understanding drug response and developing personalized cancer therapies. These three-dimensional culture systems preserve critical features of the original tumor, including cellular heterogeneity, tissue architecture, and drug sensitivity patterns that closely mirror patient clinical responses^1-3^. Individualized patient tumor organoids (IPTOs), a recently developed organoid technology for glioblastoma (GBM), have demonstrated remarkable fidelity in recapitulating tumor biology and predicting patient outcome^4^. The ability to generate multiple organoids from a single patient specimen enables parallel drug testing under controlled conditions, creating unprecedented opportunities for personalized medicine approaches. Most importantly, comparison of scRNA-seq data from treated versus untreated organoids provides deep molecular insights into drug mechanisms, response biomarkers, and resistance pathways at single-cell resolution—information that is impossible to obtain from patients directly.

However, modeling scRNA-seq perturbation data from patient-derived organoids presents significant computational challenges that extend beyond the well-recognized complexities of single-cell genomics. Particularly, predicting how individual cells will respond to perturbation demands methods that can capture both population-level shifts and cell-type-specific responses while maintaining biological interpretability. The field has witnessed rapid development of innovative computational methods specifically designed to model and predict single-cell drug responses, such as scGen^5^, Cel-1OT^6^, MOSCOT^7^, and CINEMA-0T^8^. These methods leverage deep learning architectures and mathematical frameworks to model complex cellular response patterns. However, emerging evidence suggests that simpler linear approaches may achieve comparable or even superior performance in certain contexts, raising important questions about the optimal balance between methodological sophistication and practical effectiveness^9^. This apparent paradox highlights the need for systematic benchmarking studies that can guide method selection based on specific biological contexts and research objectives.

Here, we developed DELPHAI (Deep ExplainabLe Predictive Human-organoid based AI), an AI agent specifically designed to model drug responses in patient-derived organoids using scRNA-seq perturbation data. DELPHAI integrates two analytical modules: a single-cell module for dimensionality reduction, perturbation prediction, and differential expression analysis, and a mechanistic module leveraging large language models for pathway enrichment, drug mechanism interpretation, and therapeutic recommendations.

To ensure robust method evaluation, we designed a comprehensive benchmarking framework that addresses critical gaps in current perturbation analysis assessment. Our evaluation system employs complementary metrics—mean squared error (MSE) for population-level accuracy and energy distance (E-distance) for distributional similarity—and uniquely evaluates method performance in both computational embedding spaces and biologically interpretable reconstructed gene expression spaces. This dual-space evaluation approach ensures that computational predictions maintain biological meaning and clinical relevance, addressing a key limitation in existing benchmarking studies that focus primarily on computational metrics.

We demonstrate DELPHAI’s capabilities using IPTO-derived scRNA-seq data from GBM patients treated with temozolomide, showing that the framework can accurately predict cellular responses to drug treatment, identify resistance mechanisms without prior knowledge of drug identity, and generate clinically relevant therapeutic recommendations. Through systematic benchmarking, we reveal that optimal transport combined with principal component analysis provides superior performance for capturing complex population dynamics in drug-treated organoids. Furthermore, DELPHAI’s mechanistic agent successfully predicted DNA alkylation as the primary drug mechanism and identified multiple resistance pathways, recommending targeted combination therapies that align with established clinical strategies. These results demonstrate DELPHAI’s potential to bridge computational prediction with actionable therapeutic insights, representing a significant advance toward Al-driven drug discovery and precision medicine in cancer treatment.

## Results

### DELPHAI workflow for single-cell perturbation analysis and therapeutic prediction

We developed DELPHAI (Deep ExplainabLe Predictive Human-organoid based AI), an AI agent that is specifically designed to model single-cell RNA sequencing data of patient-derived organoids under drug treatment to predict drug responses, reveal mechanistic insights and recommend personalized treatments. The complete workflow encompasses IPTO culture and drug treatment, single-cell RNA sequencing, computational perturbation prediction, and AI-driven mechanistic analysis with therapeutic recommendations (Figure 1).

**Figure 1:**
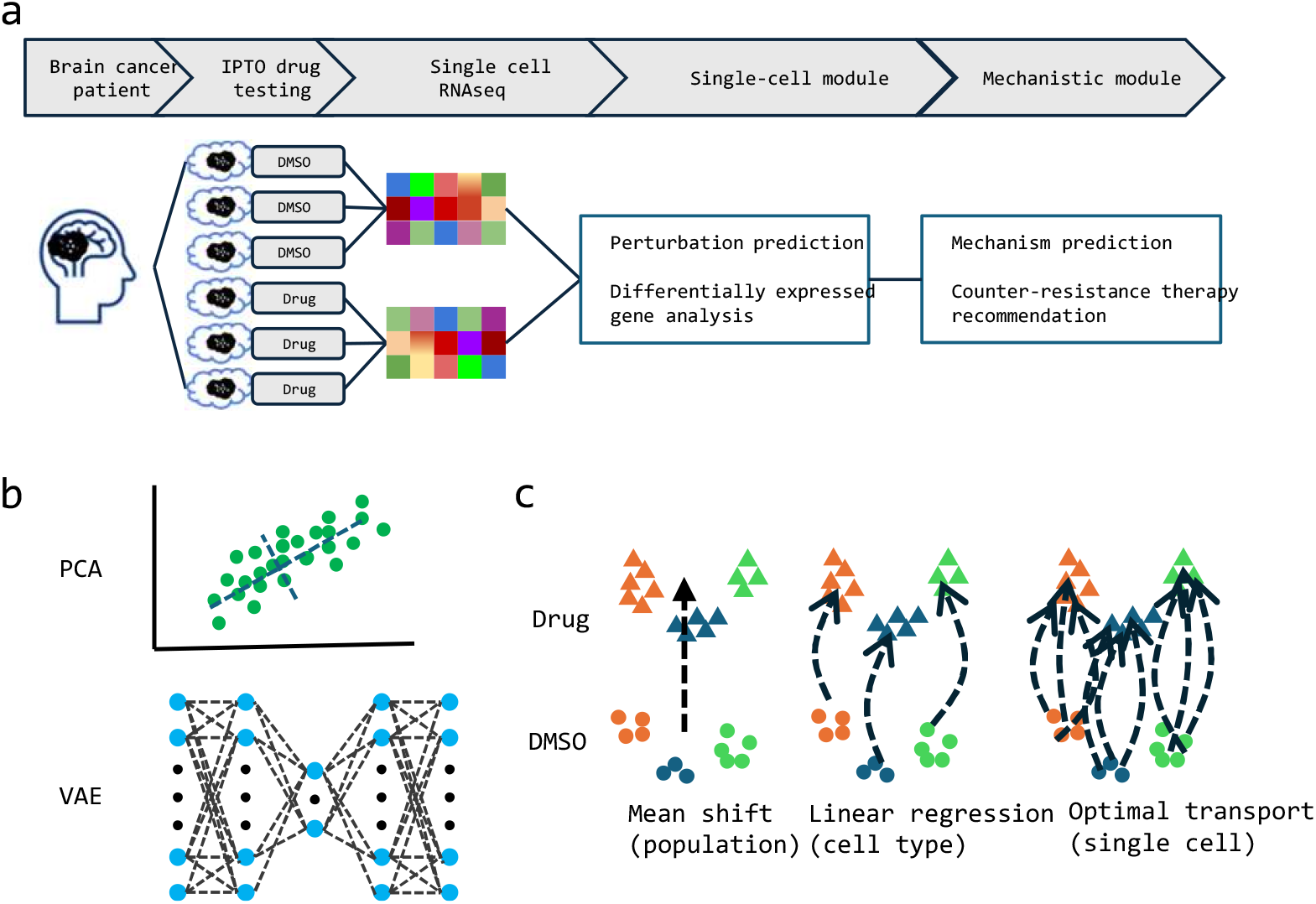
Overview and detailed methods of the workflow, **a,** workflow overview. IPTOs established from the same patient are tested against drugs and controls in parallel. Single cell RNA sequencing is performed on pooled samples from the same treatment and fed to AI agents for perturbation effect prediction, mechanism of action analysis and counter-resistance strategy recommendations, **b**, Dimensionality reduction methods benchmarked, principal component analysis and variational autoencoders, **c,** Perturbation effect prediction methods benchmarked, mean shift on the population level, linear regression per cell type and optimal transport per cell.

To demonstrate DELPHAI’s capabilities, we applied the framework to the IPTO2169 data set published previously^4^, which is scRNA-seq data of individualized patient tumor organoids (IPTOs) established from patient 2169, treated in parallel with temozolomide (TMZ) or DMSO vehicle control. IPTO2169 was shown to be sensitive to TMZ and exhibited substantial transcriptional changes in response to TMZ treatment^4^, providing an ideal test case for evaluating perturbation prediction accuracy and mechanistic studies.

Various classical and innovative algorithms have been previously developed and applied for scRNA-seq analysis and perturbation effect prediction. We first systematically benchmarked various computational methods in their performance, including principal component analysis and variational autoencoders for dimensionality reduction, and mean shift, linear regression, optimal transport for perturbation effect prediction. More details about the methods and benchmarking metrics are described in Materials and Methods.

### Benchmarking dimensionality reduction methods

Dimensionality reduction represents a critical preprocessing step that fundamentally influences downstream perturbation analysis. We systematically compared two widely-used approaches: principal component analysis (PCA) and variational autoencoders (VAE), specifically the scGen implementation optimized for single-cell perturbation studies. Data were randomly split into training (80%) and test (20%) sets. Reconstruction accuracy was evaluated by calculating the Pearson correlation between original and reconstructed gene expression values of the test data. With standard parameters as previously published, VAE achieved a median correlation of 0.269 across all genes while PCA achieved 0.121 (Figure 2a).

**Figure 2:**
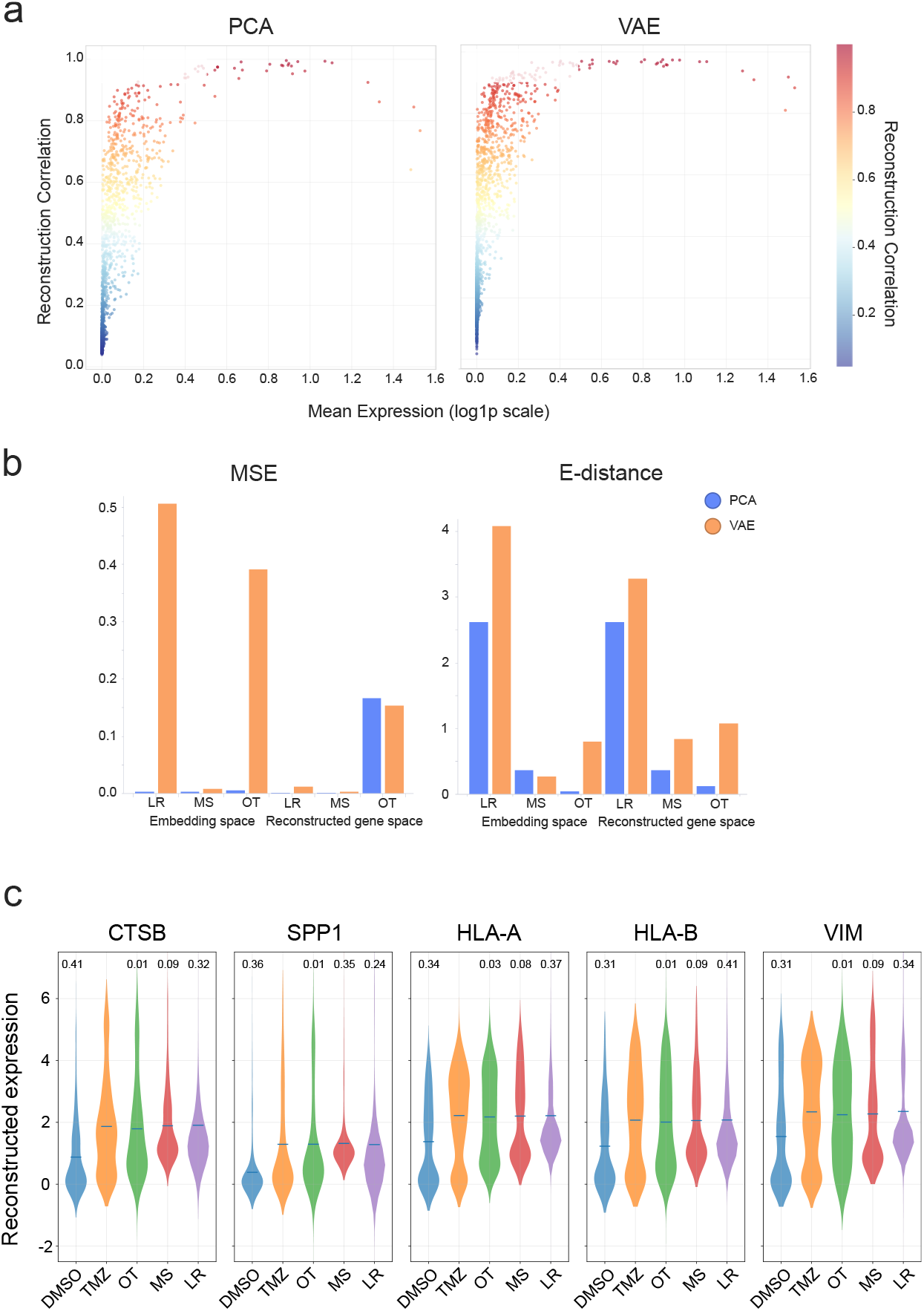
Benchmarking dimensionality reduction and perturbation effect prediction, **a,** Gene reconstruction accuracy over gene expression level for PCA (left) and VAE (right), **b**, Perturbation effect prediction quality measured by MSE (left) and E-distance (right), PCA embedding in blue and VAE embedding in orange, **c,** Top five genes with largest change due to treatment, their DMSO-treated, TMZ-treated, OT-predicted, MS-predicted and LR-predicted distributions. Each distribution is labeled with E-distance values to the TMZ-treated population.

For both methods, reconstruction accuracy correlated with mean gene expression levels. The improved reconstruction accuracy reflected the VAE’s capacity to capture non-linear relationships in gene expression data through its neural network architecture, consistent with previous reports^5,10^.

However, reconstruction accuracy alone does not determine suitability for perturbation prediction. We therefore evaluated how each dimensionality reduction method affected downstream perturbation analysis using two complementary metrics: mean squared error (MSE) for population level accuracy and energy distance (E-distance)^11^ for distributional similarity. These evaluations were performed in both embedding space and reconstructed gene expression space across three perturbation prediction methods: mean shift (MS), linear regression (LR), and optimal transport (OT).

Remarkably, despite its relatively lower reconstruction accuracy, PCA substantially outperformed VAE for perturbation effect prediction across nearly all method combinations (Figure 2b). For example, when combined with optimal transport and evaluated by E-distance in reconstructed expression space, PCA achieved an E-distance of 0.12 compared to VAE’s 1.07, when combined with mean shift and evaluated by MSE on reconstructed expression space, PCA achieved an MSE of 0.00008 compared to VAE’s 0.003. This superior performance likely reflects PCA’s preservation of global variance structure, which proves more relevant for capturing perturbation-induced population shifts than the local non-linear relationships emphasized by VAE reconstruction. Given that perturbation prediction accuracy represents the primary objective of our analysis, we selected PCA as the optimal dimensionality reduction method for subsequent analyses, reducing data to 50 principal components that captured the major axes of transcriptional variation.

### Benchmarking perturbation effect prediction methods

We next systematically evaluated three distinct approaches for predicting cellular responses to drug perturbation: mean shift at the population level, linear regression stratified by cell type, and optimal transport mapping individual cells between conditions. The mean shift approach assumes that drug effects can be captured by consistent gene expression changes across the cell population. Linear regression extends this concept by learning cell-type-specific transformation models, accounting for differential perturbation responses across cellular subpopulations. Optimal transport learns the optimal mapping between control and treatment cell distributions without assuming uniform responses across the population. Although not exhaustive, these three methods represent a good range of methods and concepts to test in our use case.

Each method was trained on the 80% training data and evaluated on the 20% test data. Evaluation using MSE revealed similar performance across methods in embedding space, with optimal transport performed substantially worse in reconstructed expression space (OT: 0.17 vs MS: 0.00008 and LR: 0.00008) (Figure 2b). Conversely, energy distance evaluation—which captures distributional rather than mean differences—revealed optimal transport’s superior performance in both embedding and reconstructed expression space (OT: 0.05 vs MS: 0.36 and LR: 2.62), demonstrating its ability to accurately capture population-level distributional changes.

This apparent contradiction reflects fundamental differences in what these metrics capture: MSE emphasizes population means, while E-distance assesses distributional similarity. Since drug perturbations often involve complex changes in cell state distributions rather than uniform shifts, E-distance provides a more biologically relevant evaluation metric. To further validate complex changes in distribution of gene expression values, we examined the five genes with the highest Edistance changes between control and TMZ-treated populations, representing those most affected by treatment in terms of distributional shifts. Optimal transport clearly outperformed both mean shift and linear regression in recapitulating the actual gene expression distributions for these treatment-responsive genes (Figure 2c), accurately capturing both the magnitude and distributional characteristics of treatment-induced changes.

### Optimal transport captures complex population dynamics

To visualize how treatment affected cell population dynamics and how predictions match ground truth, we generated UMAP projections of both control and treated populations of the 20% test data alongside predictions from each method (Figure 3a).

**Figure 3:**
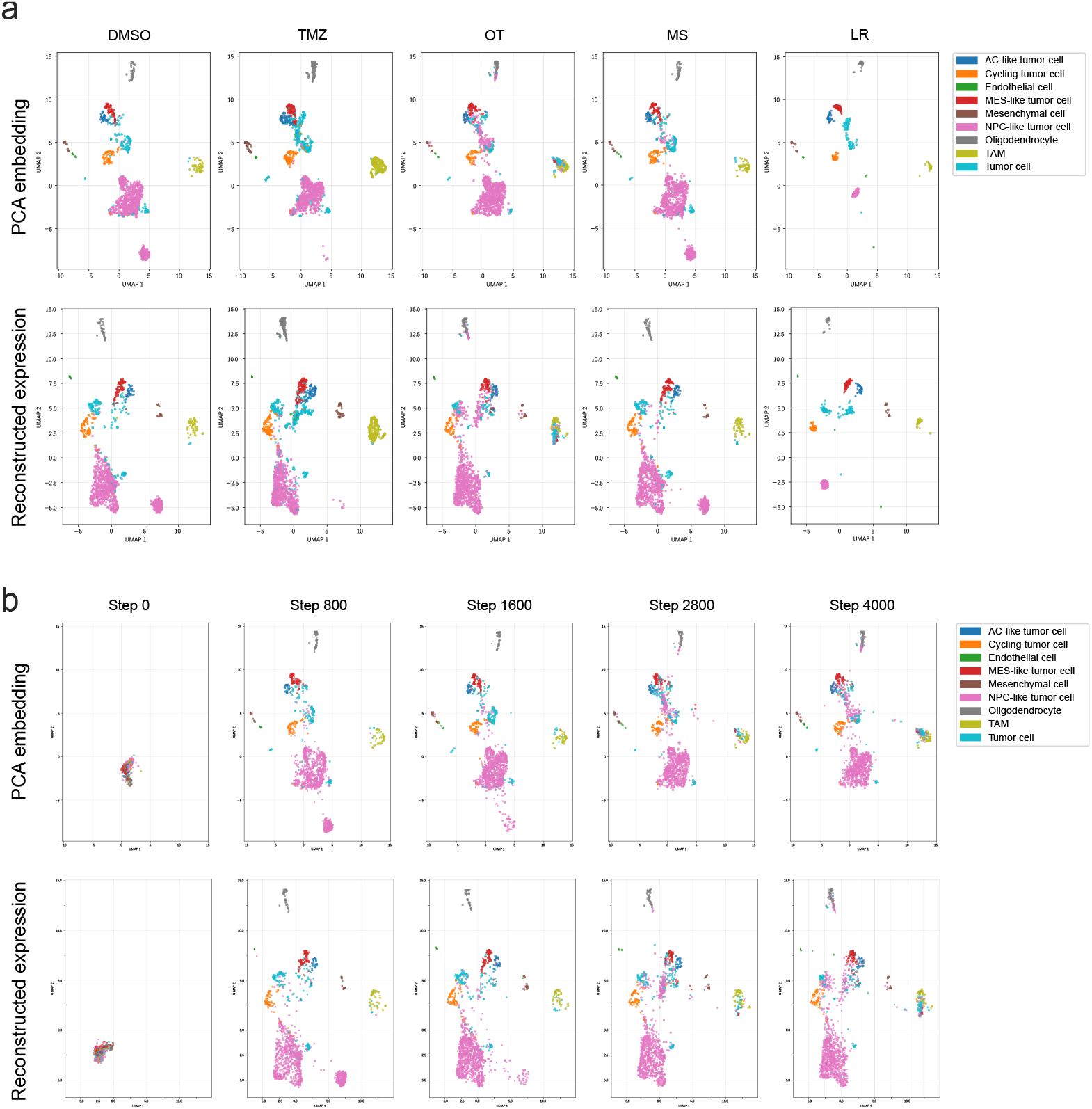
UMAP visualization of treated and predicted scRNA-seq data, **a,** DMSO-treated, TMZ-treated, OT-predicted, MS-predicted, LR-predicted cell populations on PCA embedding space and reconstructed expression space, color-coded by cell types, **b**, UMAP changes over training steps on PCA embedding space and on reconstructed expression space, color-coded by cell types.

As reported previously^4^, TMZ treatment led to substantial alterations in cellular composition: a subset of NPC-like tumor cells, represented by the cluster in pink at the bottom, was eliminated, while tumor-associated macrophages (TAMs), represented by the cluster in green to the right, increased in abundance. Notably, these changes are apparent on both embedding space and reconstructed expression space, confirming the accuracy of reconstruction. Mean shift produced uniform shifts and linear regression collapsed variations within each cell type. Predictions from both methods deviated clearly from the ground truth, not capturing the complex changes caused by drug treatment. In contrast, optimal transport accurately predicted changes in NPC-like tumor cells and TAMs, correctly identifying which cell populations would be depleted or enriched following treatment.

To understand the optimal transport training process, we analyzed predictions from checkpoint models at different training steps (Figure 3b). In both PCA embedding space and reconstructed expression space, control cells were rapidly transported (within 800 training steps) to match the general patterns of target cells, with most cell types achieving appropriate configurations. However the fine details of cell population dynamics required longer training. For example NPC-like tumor cells at the bottom started to shift upwards from step 1600, and more TAMs started to accumulate at step 2800. These results, although qualitative, demonstrate that optimal transport’s ability to learn complex, non-uniform mappings between cell populations makes it uniquely suited for capturing the heterogeneous responses that characterize drug perturbations in complex cellular systems.

### Al-driven mechanism analysis and therapeutic recommendations

With optimal transport established as the superior method for perturbation prediction, it provides a mapping between DMSO-treated and TMZ-treated cells, which in turn generated data on how single-cell-resolution expression profiles responded to the drug treatment. We hypothesized that this rich data contains the insight about response and resistance mechanisms. To test this hypothesis, we developed a customized LLM-based agent for mechanism of action (MoA) analysis, resistance characterization and counter-therapy recommendations.

To assess the agent’s ability to reveal drug mechanism insights, we challenged it to predict the drug’s MoA using only differential expression data, without revealing the drug’s identity. The agent correctly identified the primary mechanism as DNA alkylation leading to DNA damage and apoptosis, ranking “alkylating chemotherapy with immunogenic cell death” as the highest likelihood hypothesis (Table 1).

**Table 1:**
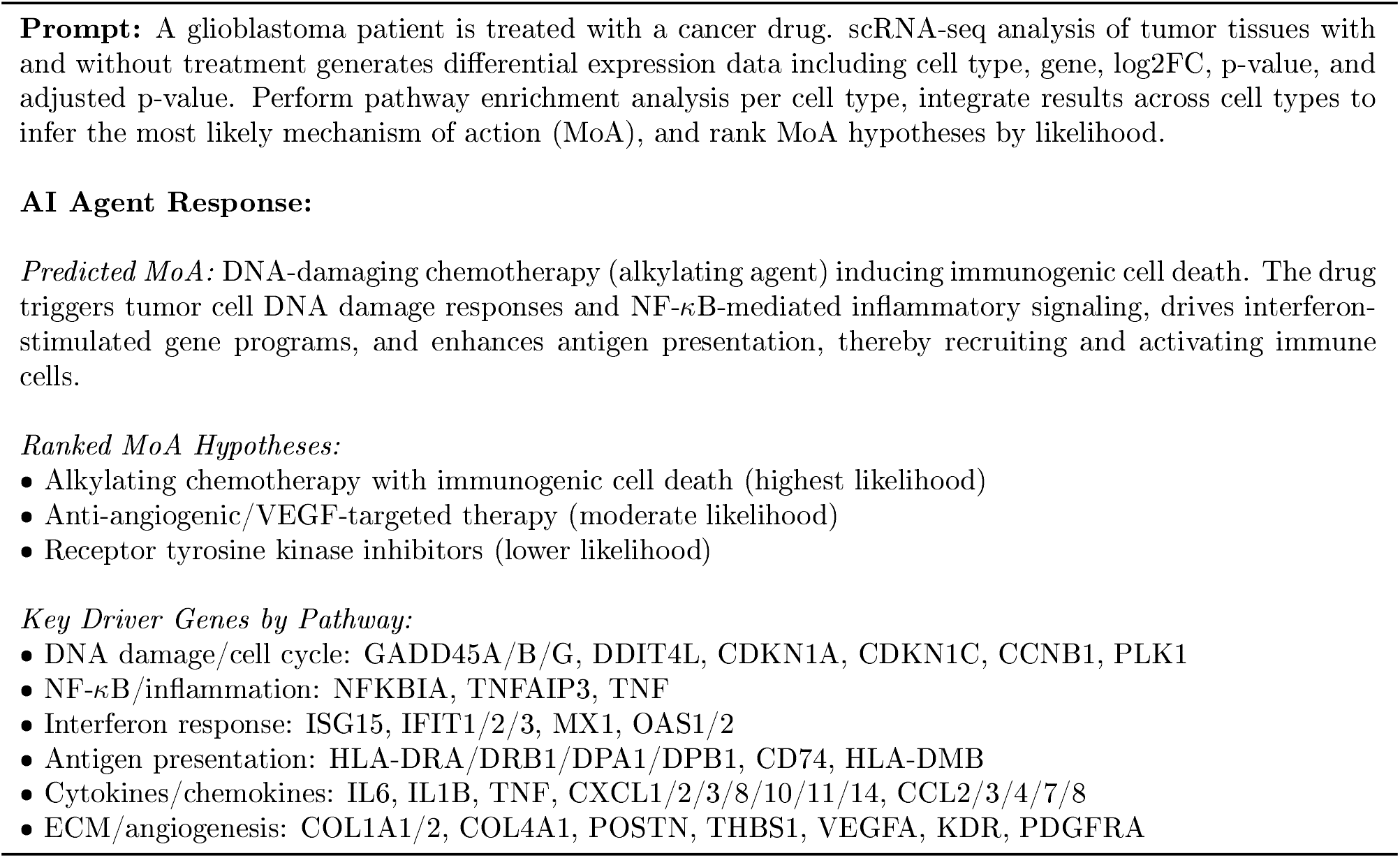
AI agent prediction of drug mechanism of action from scRNA-seq data.

This prediction aligns precisely with established knowledge that temozolomide is an alkylating agent that induces DNA damage through methylation at the 06 position of guanine^12,13^. The prediction was supported by systematic analysis of enriched pathways, including DNA damage response genes (GADD45A/B/G, CDKN1A/C), NF-kB inflammatory signaling (NFKBIA, TNFAIP3), and interferon-stimulated genes (ISG15, IFIT1/2/3), all of which are well-documented responses to TMZ treatment in glioblastoma^14,15^.

Building on this mechanistic foundation, we tasked the agent with identifying potential TMZ resistance mechanisms and recommending combination therapies. The agent systematically characterized multiple resistance pathways, prioritizing DNA damage response activation (GADD45 family upregulation, cell cycle checkpoint activation) and glioma stem cell expansion (NES, SOX2/SOX9 expression) as the most strongly supported mechanisms (Table 2).

**Table 2:**
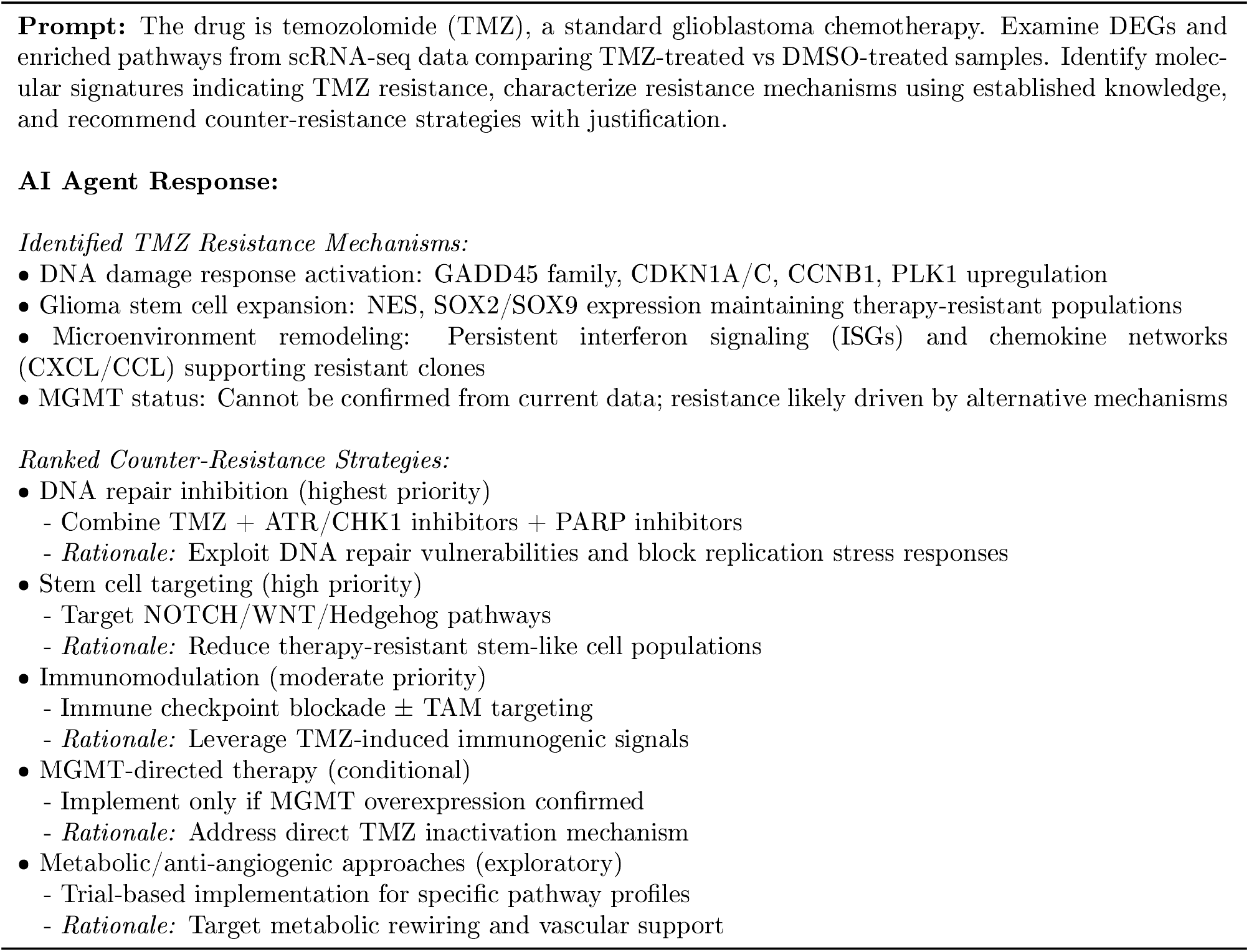
AI agent analysis of TMZ resistance mechanisms and therapeutic recommendations.

These predictions are strongly validated by clinical literature: enhanced DNA repair capacity is a major mechanism of TMZ resistance^16,17^, while glioblastoma stem cells are known to exhibit increased resistance to alkylating agents^18,19^. The agent appropriately noted that MGMT-mediated resistance could not be confirmed from transcriptional data alone, demonstrating analytical restraint consistent with the fact that MGMT resistance primarily involves protein-level activity rather than expression changes^20^.

The agent’s therapeutic recommendations were ranked by mechanistic strength and clinical feasibility. The highest priority recommendation involved combining TMZ with DNA repair inhibitors (ATR/CHK1 inhibitors plus PARP inhibitors) to exploit synthetic lethality in DNA repair pathways. This recommendation is strongly supported by ongoing clinical trials, including studies of TMZ combined with the PARP inhibitor veliparib^21^ and ATR inhibitors such as AZD6738^22^. Secondary recommendations included stem cell pathway targeting (NOTCH/WNT/Hedgehog inhibition) and immunomodulatory approaches leveraging TMZ-induced immunogenic cell death signatures. These strategies align with current therapeutic development, including clinical trials of y-secretase inhibitors targeting NOTCH signaling^23^ and combination approaches with immune checkpoint inhibitors^24^.

The systematic integration of perturbation prediction with mechanistic analysis provides a comprehensive framework for identifying resistance mechanisms and developing rational combination therapies. Importantly, the AI agent’s predictions demonstrate substantial concordance with established mechanistic understanding and ongoing clinical development, validating the approach’s ability to translate computational predictions into clinically relevant insights.

## Discussion

Our systematic benchmarking of dimensionality reduction and perturbation prediction methods reveals a fundamental disconnect between reconstruction accuracy and task-specific performance in single-cell drug response analysis. Despite VAE achieving superior gene expression reconstruction (median Pearson correlation 0.269 vs 0.121 for PCA), PCA substantially outperformed VAE in perturbation prediction across all method combinations, with improvements ranging from 2.2-fold to 8.9-fold depending on the metric and method. This finding demonstrates that task-specific evaluation is more predictive of performance than intermediate computational metrics. The superior performance of PCA likely stems from its preservation of global variance structure, which captures population-level shifts more effectively than the local non-linear relationships emphasized by VAE reconstruction.

The choice of evaluation metrics proved equally critical in revealing method performance differences. While mean squared error suggested similar performance across perturbation prediction methods, energy distance revealed optimal transport’s substantial advantages (E-distance: 0.05 vs 0.36 for mean shift and 2.62 for linear regression). This discrepancy reflects fundamental differences in what these metrics capture: MSE emphasizes population means while E-distance assesses distributional similarity. Since drug perturbations typically induce complex changes in cell state distributions rather than uniform shifts, E-distance provides a more biologically relevant evaluation framework. Our dual-space evaluation approach—assessing performance in both computational embedding space and reconstructed gene expression space—ensures that computational predictions maintain biological interpretability, addressing a critical limitation in existing benchmarking studies that focus primarily on algorithmic performance metrics.

The AI agent’s mechanism predictions demonstrated remarkable concordance with established TMZ pharmacology and clinical knowledge. The agent correctly identified DNA alkylation as the primary mechanism of action and predicted clinically validated resistance pathways, including enhanced DNA repair capacity and glioblastoma stem cell expansion. Crucially, the agent’s therapeutic recommendations—combining TMZ with PARP inhibitors and ATR/CHK1 inhibitors—align precisely with ongoing clinical trials. This validation suggests that single-cell perturbation data contains sufficient mechanistic information for AI-driven drug mechanism prediction, supporting the feasibility of automated therapeutic strategy development.

However, several limitations constrain the generalizability of our findings. The analysis relies on a single dataset from one patient and one drug, limiting conclusions about method performance across diverse tumor types, genetic backgrounds, and therapeutic agents. The heterogeneity of glioblastoma alone—spanning distinct molecular subtypes and treatment histories—necessitates validation across broader patient cohorts. Additionally, the AI agent’s reliance on existing knowledge bases may limit identification of truly novel mechanisms or unexpected drug effects. The quality of Ai-generated insights depends critically on the comprehensiveness and accuracy of training data, potentially introducing bias toward well-studied pathways and established therapeutic strategies.

Future development should prioritize several key areas to enhance clinical translation. First, systematic validation across diverse datasets, tumor types, and therapeutic agents is essential to establish method robustness and identify optimal application contexts. Second, integration of multimodal data—including imaging, genomics, proteomics, and clinical outcomes—could improve prediction accuracy and enable identification of patient-specific biomarkers. Third, incorporation of real-world evidence from electronic health records and clinical trial databases could enhance the AI agent’s therapeutic recommendations and clinical relevance. Finally, development of uncertainty quantification methods would enable clinicians to assess prediction confidence and make more informed treatment decisions.

DELPHAI represents a significant advance toward Al-driven precision medicine by demonstrating that computational analysis of patient-derived organoid data can generate clinically relevant therapeutic insights. The framework’s ability to predict drug mechanisms and resistance pathways without prior knowledge, combined with its generation of literature-validated therapeutic recommendations, suggests potential for clinical implementation. However, realizing this potential requires extensive validation, regulatory approval, and integration with existing clinical workflows. The ultimate goal is creating AI systems that not only interpret experimental data but generate testable hypotheses to advance understanding of drug action and resistance, potentially accelerating therapeutic development and improving patient outcomes in precision oncology.

## Materials and Methods

### IPTO Culture and Drug Treatment

Individualized Patient Tumor Organoids (IPTOs) were established from GBM patient 2169 and cultured following previously described protocols. Multiple IPTOs from the same patient were treated in parallel with temozolomide (TMZ) or DMSO vehicle control. Single-cell RNA sequencing was performed on pooled samples from each treatment condition. Detailed experimental procedures for IPTO establishment, culture conditions, drug treatment protocols, and scRNA-seq library preparation are described in the original IPTO publication^4^.

### Dimensionality Reduction

#### Principal Component Analysis (PCA)

PCA was applied to reduce dimensionality to 50 components, capturing major axes of transcriptional variation. The PCA model provided reconstruction capabilities through inverse transformation: 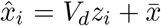, where *V_d_* contains the top 50 principal components and 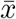 is the sample mean.

#### Variational Autoencoder

VAE was implemented using the scGen library (scgen.SCGEN) with 2-layer neural networks (512 hidden units per layer), 50-dimensional latent space, and 0.1 dropout regularization. Training used batch size 1,024, learning rate 1 × 10^−3^, maximum 1,000 epochs with early stopping, and weight decay 1 x 10^−6^.

### Perturbation Prediction

#### Mean Shift

Computed population-level differences between treatment and control conditions: Δ = *μ_t_*− *μ_c_,* then applied this shift to individual control cells: 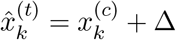.

#### Linear Regression

Learned cell-type-specific transformation matrices *W_k_* and bias vectors *b_k_*using Ridge regression for each cell type *k.* Predictions were generated as: 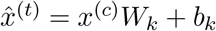 with L2 regularization parameter *λ*.

#### Optimal Transport

Implemented neural dual optimal transport using Input-Convex Neural Networks with feed-forward architectures (64 hidden units, CELU activation). Training used 2,000 optimization steps with batch size 1,024, learning rate 1 × 10^−4^, and squared Euclidean cost function. The transport map was computed as 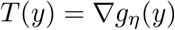 via automatic differentiation.

### Dual-Space Evaluation

Perturbation predictions were systematically evaluated in both embedding space (where models operate) and reconstructed gene expression space (where biological interpretation occurs). For PCA, gene space evaluation used inverse transformation; for VAE, forward decoder passes. Evaluation metrics included:

- **Mean Squared Error (MSE):**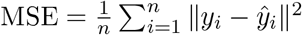
- **Energy Distance:** 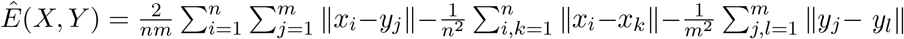

### Mechanistic Agent Implementation

A conversational AI agent was implemented using AnythingLLM^25^, an open-source platform that interfaces with large language models including ChatGPT^26^. The agent was designed with two specialized analytical modules: (1) A mechanism of action module that predicted drug mechanisms from differential expression data without prior drug knowledge, performing automated pathway enrichment analysis and ranking mechanistic hypotheses by supporting evidence; (2) A resistance analysis module that systematically identified potential resistance mechanisms and recommended targeted combination therapies based on established molecular pathways and clinical literature. Agent prompts were engineered to be specific and instructive, with responses edited for clarity and conciseness in presentation.

## Acknowledgements

aiPTO TechBio receives AWS Cloud credits through the NVIDIA Inception program and AWS Startups program. H.L. is supported by the Deutsche Forschungsgemeinschaft (DFG) (project 404521405-CRC1389 UNITE Glioblastoma A08N.), the Carl Zeiss foundation (AI-Care), the DKTK Joint Funding (AIM2GO).

